# Ultrasonic Cigarettes: Chemicals and Cytotoxicity are Similar to Heated-Coil Pod-Style Electronic Cigarettes

**DOI:** 10.1101/2024.02.28.582598

**Authors:** Esther E. Omaiye, Wentai Luo, Kevin J. McWhirter, James F. Pankow, Prue Talbot

## Abstract

Our purpose was to test the hypothesis that ultrasonic cigarettes (u-cigarettes), which operate at relatively low temperatures, produce aerosols that are less harmful than heated-coil pod-style electronic cigarettes (e-cigarettes). The major chemicals in SURGE u-cigarette fluids and aerosols were quantified, their cytotoxicity and cellular effects were assessed, and a Margin of Exposure risk assessment was performed on chemicals in SURGE fluids. Four SURGE u-cigarette flavor variants (“Watermelon Ice,” “Blueberry Ice,” “Green Mint,” and “Polar Mint”) were evaluated. Flavor chemicals were quantified in fluids and aerosols using gas chromatography/mass spectrometry. Cytotoxicity and cell dynamics were assessed using the MTT assay, live-cell imaging, and fluorescent microscopy. WS-23 (a coolant) and total flavor chemical concentrations in SURGE were similar to e-cigarettes, while SURGE nicotine concentrations (13 - 19 mg/mL) were lower than many 4^th^ generation e-cigarettes. Transfer efficiencies of dominant chemicals to aerosols in SURGE ranged from 44 - 100%. SURGE fluids and aerosols had four dominant flavor chemicals (> 1 mg/mL). Toxic aldehydes were usually higher in SURGE aerosols than in SURGE fluids. SURGE fluids and aerosols had aldehyde concentrations significantly higher than pod-style e-cigarettes. Chemical constituents, solvent ratios, and aldehydes varied among SURGE flavor variants. SURGE fluids and aerosols inhibited cell growth and mitochondrial reductases, produced attenuated and round cells, and depolymerized actin filaments, effects that depended on pod flavor, chemical constituents, and concentration. The MOEs for nicotine, WS-23, and propylene glycol were < 100 based on consumption of 1 - 2 SURGE cigarettes/day. Replacing the heating coil with a sonicator did not eliminate chemicals, including aldehydes, in aerosols or diminish toxicity in comparisons between SURGE and other pod products. The high concentrations of nicotine, WS-23, flavor chemicals, and aldehydes and the cytotoxicity of SURGE aerosols do not support the hypothesis that aerosols from u-cigarettes are less harmful than those from e-cigarettes.

## INTRODUCTION

During the last decade, there has been exponential growth in the production, distribution, and use of electronic cigarettes (e-cigarettes) by never-before users of tobacco products and smokers trying to quit.^1–5^ During this time, e-cigarettes have continually evolved, and new products with modified designs and novel chemical ingredients have had a strong appeal, especially among adolescents and young adults.^6–9^ There are currently four generations of e-cigarettes based on atomizer design and fluid composition.^10–12^ Fourth-generation e-cigarette fluids (e-liquids) contain nicotine (freebase or salt base), solvents (mainly propylene glycol and glycerol), synthetic coolants (mainly WS-23 and WS-3), and a wide range of characterizing and non-characterizing flavor chemicals.^13–17^ The cellular, physiological, and potential health effects of individual constituents and their mixtures have been studied using in vitro models, experimental animals, and humans.^18–23^ These studies have contributed to regulatory policies intended to limit the sale of e-cigarettes, especially to adolescents and young adults, and to require premarket approval by the FDA.^24–27^ Local and Federal policies have caused manufacturers to innovate around regulations by developing new designs, such as disposable 4th generation products and formulations, such as the use of synthetic nicotine.^28–30^

Ultrasonic cigarettes (u-cigarettes) are interesting new tobacco products that aerosolize flavored fluids using ultrasonic waves.^31^ While initial entries into this market did not perform well, SURGE u-cigarettes, manufactured by Innokin, are gaining traction in both online reviews and sales^32,33^ and are being marketed with little information on their chemical composition and safety. Rather than heating a coil, SURGE products aerosolize a fluid containing nicotine, flavor chemicals, and solvents at very high frequencies, which may produce fewer harmful reaction products than traditional e-cigarettes.^34^ Current research on u-cigarettes is limited to a single study showing that u-cigarettes impaired flow-mediated dilation of arteries in rats with low serum nicotine levels compared to IQOS and e-cigarettes.^35^

SURGE makes several claims on its website, such as “Ultrasonic technology allows SURGE to operate at lower temperatures than traditional devices. …This allows SURGE to maintain the chemical stability of e-liquid and reduce the emission of potential toxins to levels even lower than traditional vaping devices. SURGE produces the purest vapor of any device on the market.” ^36^ However, these claims have not yet been substantiated by an independent laboratory. To provide data on this new tobacco product, our study quantified the chemicals in SURGE fluids and aerosols, examined their toxicity, and compared SURGE data to other pod-based e-cigarettes and refill fluids.

## MATERIALS AND METHODS

### Materials

For gas chromatography/mass spectrometry (GC/MS) analysis, isopropyl alcohol (IPA) was purchased from Fisher Scientific (Chino, CA). For cell culture and cell-based assays, Dulbecco’s phosphate buffered saline (DPBS) and dimethyl sulfoxide (DMSO) were purchased from Fisher Scientific (Chino, CA). BEAS-2B cells were obtained from American Type Cell Culture (ATCC, USA). Bronchial epithelial basal medium (BEBM) and supplements were purchased from Lonza (Walkersville, MD). Collagen (30 mg/mL), bovine serum albumin (BSA, 10 mg/mL) fibronectin (10 mg/mL), poly-vinyl-pyrrolidone (PVP), and MTT reagent (3-(4,5-dimethylthiazol-2-yl)-2,5-diphenyltetrazolium bromide) were purchased from Sigma-Aldrich (St Louis, MO). Phalloidin-iFluor 594 was purchased from Abcam, Cambridge, United Kingdom.

Sample Acquisition. Rechargeable SURGE (Shenzhen Innokin Technology Co. Ltd) u-cigarettes were purchased online from www.myvaporstore.com. SURGE u-cigarettes have a 700 mAh rechargeable battery permitting a 1.5A current flow that produces an aerosol at 5V (volts)/7.5W (watts). “Blueberry Ice,” “Watermelon Ice,” “Green Mint,” and “Polar Mint” flavors were purchased in pre-filled pods containing 1.2 mL of fluid.

### Aerosol Sample Preparation, Production, and Capture

Flavored pods were primed for aerosolization by taking three puffs and the weights before aerosol production. The generated aerosol was captured in isopropyl alcohol (IPA) for chemical analysis or in basal culture medium for cell analysis.^13^ Two 125 mL impingers set up at room temperature were connected to a Cole-Parmer Masterflex L/S peristaltic pump, and pods were puffed using a 4.3 s puff duration^37^ with inter-puff intervals of 60 seconds and an airflow rate of 10 mL/s. The fluid level was monitored to avoid vaping beyond ¾ of the pod’s capacity to avoid “dry puffing.” Pods were weighed before and after aerosol production to collect at least 10 mg in 30 mL of IPA for GC/MS. Aerosol solutions were stored at −20 °C and analyzed within 2 days. The number of puffs taken to achieve > 10 mg in weight was variable; “Blueberry Ice (120 puffs),” “Watermelon Ice (90 puffs),” “Green Mint (60 puffs),” and “Polar Mint (90 puffs).”

For cell-based assays, 6 total puff equivalents or TPEs (1 TPE = 1 puff/mL of culture medium) of aerosol solution were collected in 25 mL of BEAS-2B basal medium, supplemented after aerosol production to obtain a complete growth medium. The complete medium was filtered using a 0.2 µm filter, and aliquots were stored at -80 °C until testing. Aerosols were tested at concentrations ranging from 0.02 - 6 TPE. The TPE concentrations were converted to percentages of the pod fluid by considering the pod weight difference before and after aerosol collection and determining the weight of the fluid consumed. The weight (grams) of fluid consumed/puff of aerosol was calculated, and the density of the pod fluid was determined. Then, the grams/puff were converted to milliliters using the density values. Finally, the percent for concentrations used in the aerosol cell-based assays was determined according to the equation: (Np x Vp)/Vm where Np is the number of puffs, Vp is the volume of 1 puff, and Vm is the volume of the medium.

For aldehyde quantification in condensates, 120 puffs of aerosols were collected in two 30 mL mini impingers set up in an acetone dry ice bath with a temperature of -78 °C. After the puffing section was completed, the mini impingers were allowed to warm up, and condensate material was collected and stored at -20 °C for 1-2 days before analysis.

### Quantification of Chemicals Using GC/MS

Unvaped fluid collected from pods was analyzed using previously described GC/MS methods.^13,38^ 50 μL of each sample were dissolved in 0.95 mL of IPA and shipped overnight on Ice to Portland State University. Before analysis, a 20 μL aliquot of internal standard solution (2000 ng/μL of 1,2,3-trichlorobenzene dissolved in IPA) was added to each diluted sample. Analyses were performed for 178 flavor-related target analytes, two synthetic coolants, and nicotine with an Agilent 5975C GC/MS system (Santa Clara, CA). The GC column was a Restek Rxi-624Sil MS column (Bellefonte, PA) (30m long, 0.25 mm id, and 1.4 μm film thickness). A 1.0 μL aliquot of diluted sample was injected into the GC at 235 °C with a 10:1 split. The GC temperature program for analyses was: 40 °C hold for 2 min, 10 °C/min to 100 °C, then 12 °C/min to 280 °C and hold for 8 min at 280 °C, then 20 °C/min to 230 °C. The MS was operated in electron impact ionization mode at 70 eV in positive ion mode. The ion source temperature was 220 °C, and the quadrupole temperature was 150 °C. The scan range was 34 to 400 amu. Each of the 181 (178 flavor chemicals, two synthetic coolants, and nicotine) target analytes were quantitated using authentic standard materials using internal-standard-based calibration procedures described elsewhere.^38^

### Analysis of Aldehyde-Related Reaction Products Using GC/MS

Twelve (12) target analytes were investigated in pod fluids and aerosol condensates. 1 mL of HPLC grade water, e-fluid sample (50 µL), internal standard solution (20 μL at 2 mg/mL of 2′,4′,5′-Trifluoroacetophenone dissolved in acetonitrile/water, 50/50), and derivatization solution (1 mL of 12 mg/ mL PFBHA in PH=4 citric acid buffer) were added into a 5 mL Reacti-vial and vortexed for 3 times at 10 seconds each, then covered with aluminum foil and left at room temperature for 24 hours. After 24 hours, 4 drops of 40% H2SO4 and 1 mL dichloromethane (DCM) were added to the mixture solution, vortexed 3 times at 10 seconds each, and placed at room temperature for a 30-minute extraction. After 30 mins, the vial was centrifuged for 3 min, and the bottom layer was collected into a test tube with 80 mg Na2SO4. The DCM solution was then transferred into an autosampler vial for GCMS analysis. The GC/MS system was an Agilent 5975C (Santa Clara, CA). A Restek Rxi-624Sil MS GC column (30 m, 0.25 mm id, and 1.4 µm film thickness) (Bellefonte, PA) was used for the separation. The GC oven program was: 70 °C hold for 2 min; 10 °C/min to 100 °C; 5 °C/min to 250 °C; then 10 °C/min to 280 °C hold for 10 min at 280 °C; then 25 °C/min to 230 °C. The MS was operated in electron impact ionization mode at 70 eV in positive ion mode. The ion source temperature was 250 °C, and the scan range was 50 to 500 amu.

### Estimation of Non-Target Chemicals

The total ion chromatogram (TIC) response factor of the internal standard in each data set was used to estimate the ng/µL concentration of each non-target chemical from their TIC peak areas based on the assumption that each chemical’s response factor (peak area per ng) was similar to that of the internal standard (1,2,3-trichlorobenzene). While fluid samples were diluted by a factor of 20, aerosol samples were not diluted. The mass concentration is expressed as µg/mL of undiluted e-fluid (for e-fluid) or condensate (for aerosol) samples. The estimation limit was ∼0.001 mg/mL.

### Human Bronchial Epithelial Cell (BEAS-2B) Culture and Cellular Assays

BEAS-2B cells (passages 30 - 34) were cultured in BEGM (bronchial epithelial growth medium) supplemented with 2 ml of bovine pituitary extract and 0.5 ml each of insulin, hydrocortisone, retinoic acid, transferrin, triiodothyronine, epinephrine, and human recombinant epidermal growth factor.^39^ Nunc T-25 tissue culture flasks were coated overnight with BEBM fortified with collagen (30 mg/mL), bovine serum albumin (BSA, 10 mg/mL), and fibronectin (10 mg/mL) before culturing. Cells were maintained at 30 - 90% confluence at 37o C in a humidified incubator with 5% carbon dioxide. For sub-culturing, cells were harvested using DPBS for washing and incubated with 1.5 ml of 0.25% trypsin EDTA/DPBS and PVP for 3–4 mins at 37°C to allow detachment. Cells were counted using a hemocytometer and cultured in T-25 flasks at 75,000 cells/flask. The medium was replaced every other day. For in-vitro assays, cells were cultured and harvested at 80-90% confluency using protocols previously described.^39^

### MTT Cytotoxicity Assays

The effect of u-cigarette pod fluids (0.03 - 10%), aerosol condensates (0.03 - 10%), and aerosol solutions (0.02 - 6 TPE, which is equivalent to 0.004 -1.8 % of the fluid) on mitochondrial reductase activity was evaluated using treatments serially diluted in culture medium. Negative controls (0%) were placed next to the highest and lowest concentrations to check for a vapor effect from the treatments.^40^ BEAS-2B cells were seeded at 5,000 cells/well in pre-coated 96-well plates and allowed to attach for 48 hours, after which cells were exposed to treatments for 24 hours before the MTT assays. The MTT assay measures the activity of mitochondrial reductases, which convert water-soluble MTT salt to a formazan that accumulates in viable cells. After treatment, 20 µL of MTT reagent dissolved in 5 mg/mL of DPBS were added to wells and incubated for 2 hours at 37°C. Solutions were removed from wells, and 100 µl of DMSO was added to each well and gently mixed on a shaker to solubilize formazan crystals. Absorbance readings of control and treated wells were taken against a DMSO blank at 570 nm using an Epoch microplate reader (Biotek, Winooski, VT).

### Cell Growth (Area) and Morphology Assays

Time-lapse imaging was performed over 48 hours using 10x and 20x phase contrast objectives in a BioStation CT with automatic Z-focus.^41^ BEAS-2B cells were harvested and plated at 21,000 cells/well in pre-coated 24-well plates and allowed to attach for 48 hours. After attachment, BEAS-2B cells were treated with 0.6 and 6 TPE aerosol concentrations. Images were taken every 4 hours for 48 hours to collect time-lapse data for cell growth as a function of cell area (10x) and morphology (20x) analysis. BEAS-2B growth and morphology were compared in control and treated groups using CL Quant software (DR Vision, Seattle, WA).^41,42^ Data from the treated groups were normalized to untreated controls.

### Effects of Aerosol on Actin Filaments using Phalloidin-iFluor 594

The effects of aerosol treatment on actin filaments were investigated using phalloidin-iFluor 594. BEAS-2B cells were seeded at 5,000 cells/well in pre-coated 8-well Ibidi™ chamber slides (Ibidi®, Grafelfing, Germany) and attached for 48 hours before treatment with aerosols for 24 hours. After treatment, cells were rinsed with PBS (+) and fixed with 4% paraformaldehyde for 10 mins. Fixed cells were washed twice with PBS (+), labeled with phalloidin-iFluor 594 for 60 mins, washed twice with PBS (+), and incubated with a mounting shield at room temperature for 15 minutes. Cells were imaged with a Nikon Ti Eclipse inverted microscope (Nikon Instruments, Melville, NY, USA) using a 60X objective. Image processing was done using Nikon Elements.

### Data and Statistical Analyses

All statistical analyses were performed in Prism (GraphPad, San Diego, CA). Comparison of means for chemicals in different products was done using a one-way analysis of variance (ANOVA) with Dunnett’s post hoc test or an unpaired t-test on the raw data. When the conditions for the statistical analysis (homogeneity of variance and normal distribution) were not satisfied, the data were transformed using Y = Log(Y) and subjected to a one-way ANOVA. A Welch’s correction was used when t-tests were performed on the transformed data. (Figure 2f). Outliers were identified and removed from the statistical analyses using the robust regression and outlier (ROUT) removal function with the Q = 1%. In the live cell imaging assay, a two-way ANOVA with Dunnett’s post hoc test was used to compare time and concentration to the untreated control.

## RESULTS

### Propylene Glycol (PG) and Glycerol (G) Concentrations in SURGE Fluids and Aerosols

All SURGE fluids and aerosols contained PG and G. The concentration of PG and G in unvaped fluids was about 200 mg/mL and 400 mg/mL, respectively (Figures 1 A and B). The sum of both solvents ranged from 613 - 656 mg/mL in fluids and 318 - 667 mg/mL in aerosols. The ratio of PG/G in fluids was approximately 40/60 in mint flavors and 30/70 in ice flavors (Figure 1A and B). Transfer efficiencies ranged between 50% - 114% for propylene glycol and 44% - 108% for glycerol, with the lowest transfers in “Watermelon Ice” (Figure 1A and B).

**Figure 1.**
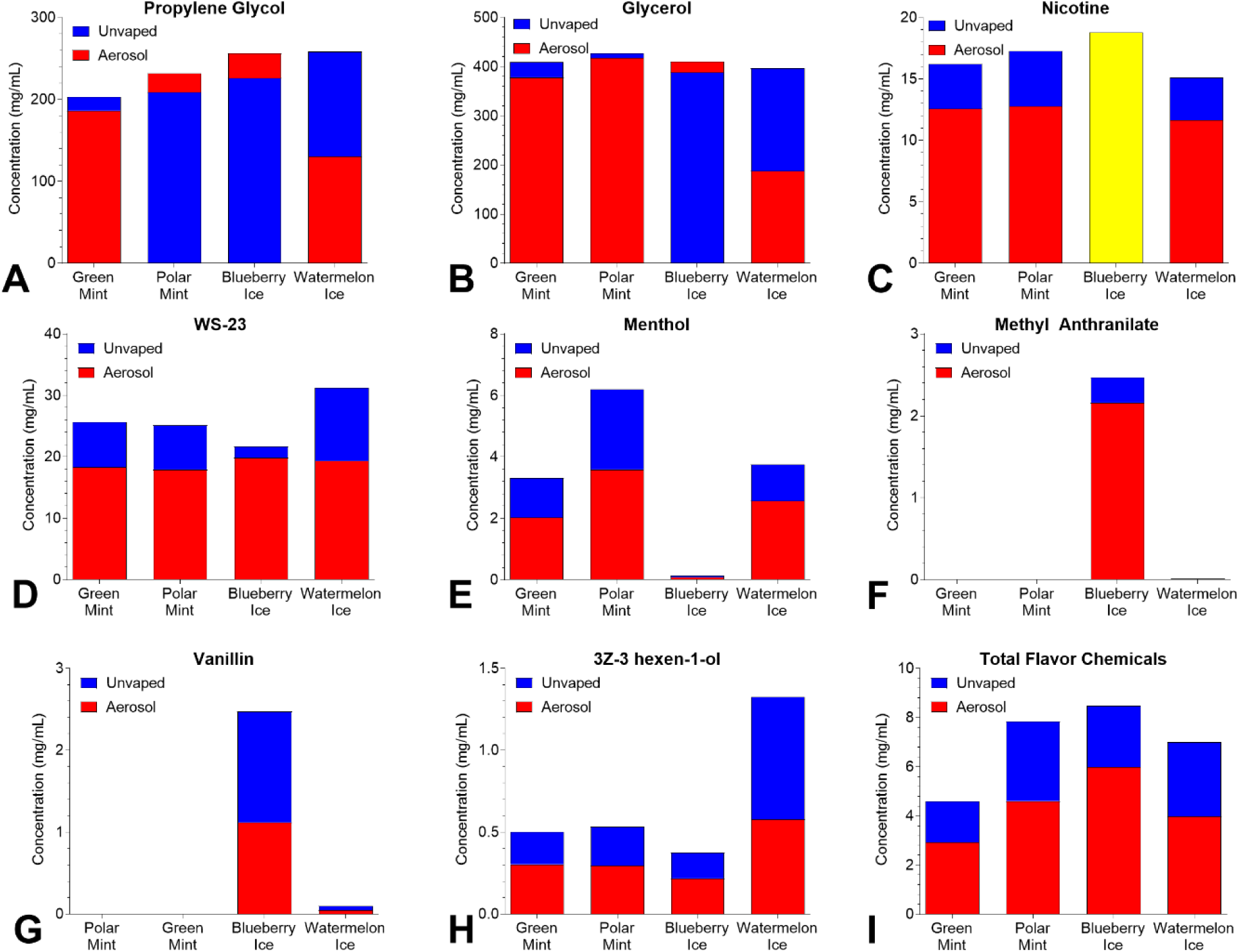
Concentrations of dominant chemicals (> 1mg/mL) in unvaped fluids and aerosols from SURGE u-cigarettes. (A) Propylene glycol, (B) Glycerol, (C) Nicotine, (D) WS-23, (E - H) Menthol, methyl anthranilate, vanillin, and (3Z)-3-hexen-1-ol, concentrations were > 1mg/mL in the fluid of at least 1 product, and (I) “Total Flavor Chemicals.” Flavor chemicals with concentrations between 0.01 - 0.99 mg/mL are in Figure S2. The yellow bar indicates that concentrations were the same in fluid and aerosol. Data show single measurements from one pod and do not include values below the LOQ.

### Nicotine and WS-23 in SURGE Fluids and Aerosols

Nicotine concentrations in fluid samples ranged from 15.1 - 18.8 mg/mL, with “Blueberry Ice” containing equal amounts in both fluid and aerosol samples (Figure 1C). Transfer efficiencies for nicotine from the fluid into the aerosol varied between 74 - 100% depending on the flavor variant, with a 100% transfer efficiency in “Blueberry Ice.”

WS-3 was not detected in any of the SURGE pods; however, WS-23 was in both “mint” and “ice” flavors at concentrations ranging from 21.7 – 31.3 mg/mL in unvaped fluids and 17.9 – 19.9 mg/mL in aerosols with transfer efficiencies ranging from 62 – 92 % (Figure 1D).

### Concentrations of Flavor Chemicals in SURGE Fluids and Aerosols

Of the 180 flavor chemicals on our target list, 74 (41%) were identified in SURGE fluids and aerosols (Figure 1E-I, Figure S1, and Table S1). Chemicals below the LOQ with estimated concentrations are listed in Table S1. 46 of 74 chemicals above the LOQ (0.01 mg/mL for fluids and 0.02 mg/mL for aerosols) are shown in Figure 1 E-I and Figure S1.

Four dominant flavor chemicals (>1 mg/mL) were identified and were relatively low in concentration (range = 1.4 mg/mL for 3Z-3 hexen-1-ol to 6 mg/mL for menthol) in (Figure 1 E – H). While menthol and 3Z-3-hexen-1-ol were dominant in “Watermelon Ice” with transfer efficiencies of 68% and 44%, respectively, methyl anthranilate and vanillin were dominant in “Blueberry Ice” with transfer efficiencies of 45% and 87%, respectively (Figure 1 E - H). The total concentration of flavor chemicals ranged from 4.6 - 8.5 mg/mL in unvaped fluids and 2.9 – 6.0 mg/mL in aerosols, with transfer efficiencies ranging from 57% – 71 %. (Figure 1I).

### Chemical Concentrations in SURGE Compared to E-cigarette Products

Nicotine, WS-23, and Total Flavor Chemical concentrations in SURGE were compared to previously published data from JUUL, PUFF, and LiQua products (Figure 2 A-F).^13,14,39^ Nicotine concentrations were significantly lower in unvaped SURGE fluids than in JUUL and PUFF fluids but significantly higher than in LiQua refill fluids (Figure 2A). Aerosol nicotine was likewise higher in JUUL than in SURGE (Figure 2D). WS-23, a synthetic coolant, was significantly higher in SURGE fluids than in JUUL and LiQua (Figure 2B), and WS-23 was significantly higher in the aerosol from SURGE than from JUUL (Figure 2E). While WS-23 concentrations in Puff were highly variable with flavor variants, all surge products had similar WS-23 concentrations. The concentration of Total Flavor Chemicals in SURGE fluids and aerosols was not significantly different from JUUL, PUFF, or LiQua (Figure 2C). Total flavor chemical concentration in the SURGE aerosols was not significantly higher than in JUUL (Figure 2F).

**Figure 2.**
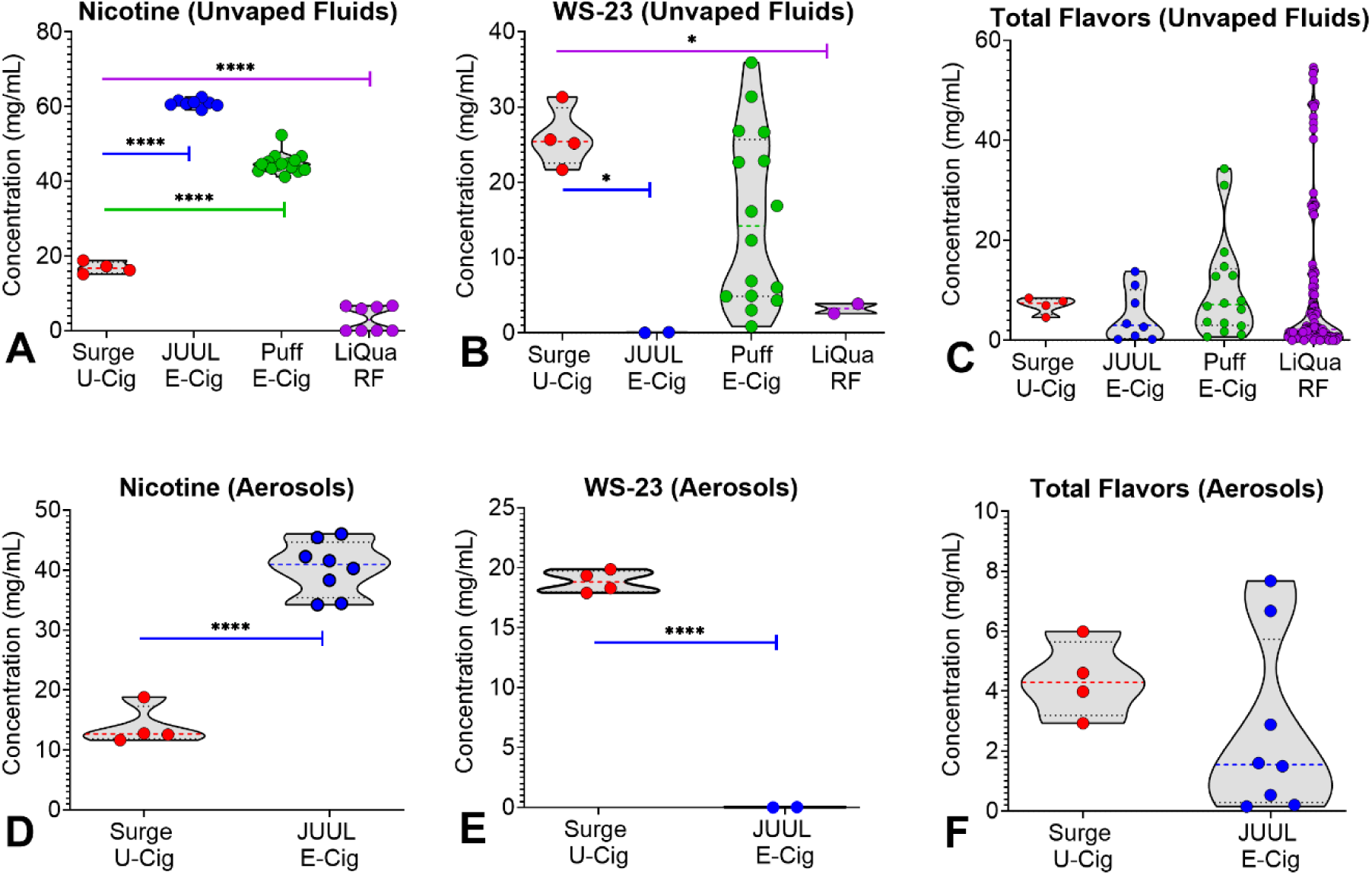
Concentrations of chemicals in SURGE, JUUL, Puff, and LiQua products. (A) Nicotine in unvaped fluids, (B) WS-23 in unvaped fluids, (C) “Total Flavor Chemicals” in unvaped fluids, (D) Nicotine in aerosols, (E) WS-23 in aerosols, and (F) “Total Flavor Chemicals” in aerosols. U-Cig = u-cigarette, E-Cig = e-cigarettes, RF = refill fluids. * = p < 0.05, ** = p < 0.01, *** = p < 0.001, **** = p < 0.0001. E-cigarette data are taken from Omaiye et al. 2019, Omaiye et al. 2020, and Omaiye et al.2022.

### Aldehydes in SURGE Fluids and Aerosols

Seven of 12 target aldehydes were detected in SURGE fluids and aerosols (Figure 3A-H). All seven aldehydes were present in the unvaped fluids from each product at concentrations ranging from 2.4 µg/mL for dihydroxyacetone in “Polar Mint” to 53.4 µg/mL for methylglyoxal in “Polar Mint.” Values below the LOQ (20 µg/mL) are estimates. Except for acetaldehyde (3 - 6 µg/mL) and acrolein (2 - 6 µg/mL), concentrations were higher in the aerosols than in the unvaped fluids. Methylglyoxal (37 - 125 µg/mL) reached the highest concentration in the aerosols with “Watermelon Ice” having 125 µg/mL (Figure 3H). The total aldehyde concentration in the aerosols ranged from 99 to 354 µg/mL, with “Green Mint” being the highest (Figure H).

**Figure 3.**
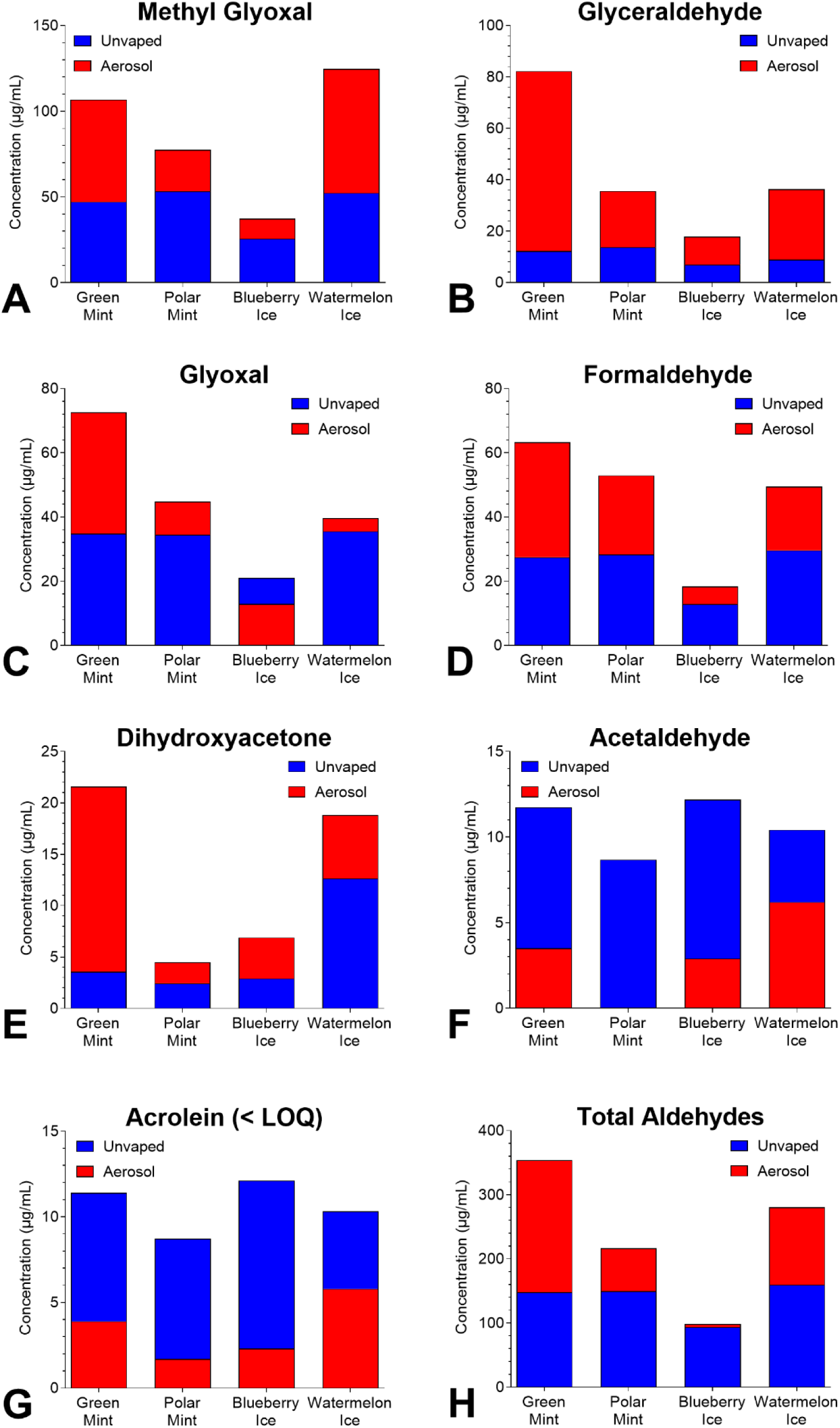
Concentrations of aldehydes in SURGE u-cigarettes. (A) Methylglyoxal, (B) Glyoxal, (C) Glyceraldehyde, (D) Formaldehyde, (E) Dihydroxyacetone, (F) Acetaldehyde, (G) Acrolein, and (H) Total Aldehydes. Acrolein was < LOQ in all samples but was higher in unvaped fluids than in aerosols. Data show single measurements from one pod and exclude values below the LOQ (except for acrolein).

### Aldehyde Concentrations in SURGE Compared to E-cigarette Products

Aldehyde concentrations in SURGE fluids and aerosols were compared to “JUUL” and “Other Brands,” which included Ezzy Oval, Hyde, Puff Bar, and SEA (Figure 4).^43–58^ Methylglyoxal and glyoxal were significantly higher in SURGE fluids and aerosols than in aerosols from JUUL and “Other Brands” (Figures 4 A, C). Glyceraldehyde was significantly higher in SURGE aerosols than in unvaped fluids (Figure 4B). Formaldehyde in SURGE fluids was within the range reported for other pod-based e-cigarette fluids but significantly higher in SURGE aerosols than in “JUUL” and “Other Brands” (Figure 4D). Although dihydroxyacetone was higher in SURGE aerosols than fluids, the difference was not statistically significant (Figure 4E). Acetaldehyde in SURGE fluids was within the range of e-cigarettes and not statistically different than levels in other pod-based e-cigarette fluids; however, it was significantly higher in SURGE aerosols than in JUUL and other e-cigarette aerosols (Figure 4F). Acrolein concentrations were under the reported limit and were higher in SURGE fluids and aerosols than in JUUL and “Other Brands” (Figure 4G). While present in JUUL and Other Aerosols, propionaldehyde and crotonaldehyde were not found in SURGE aerosols (Figure 4H).

**Figure 4:**
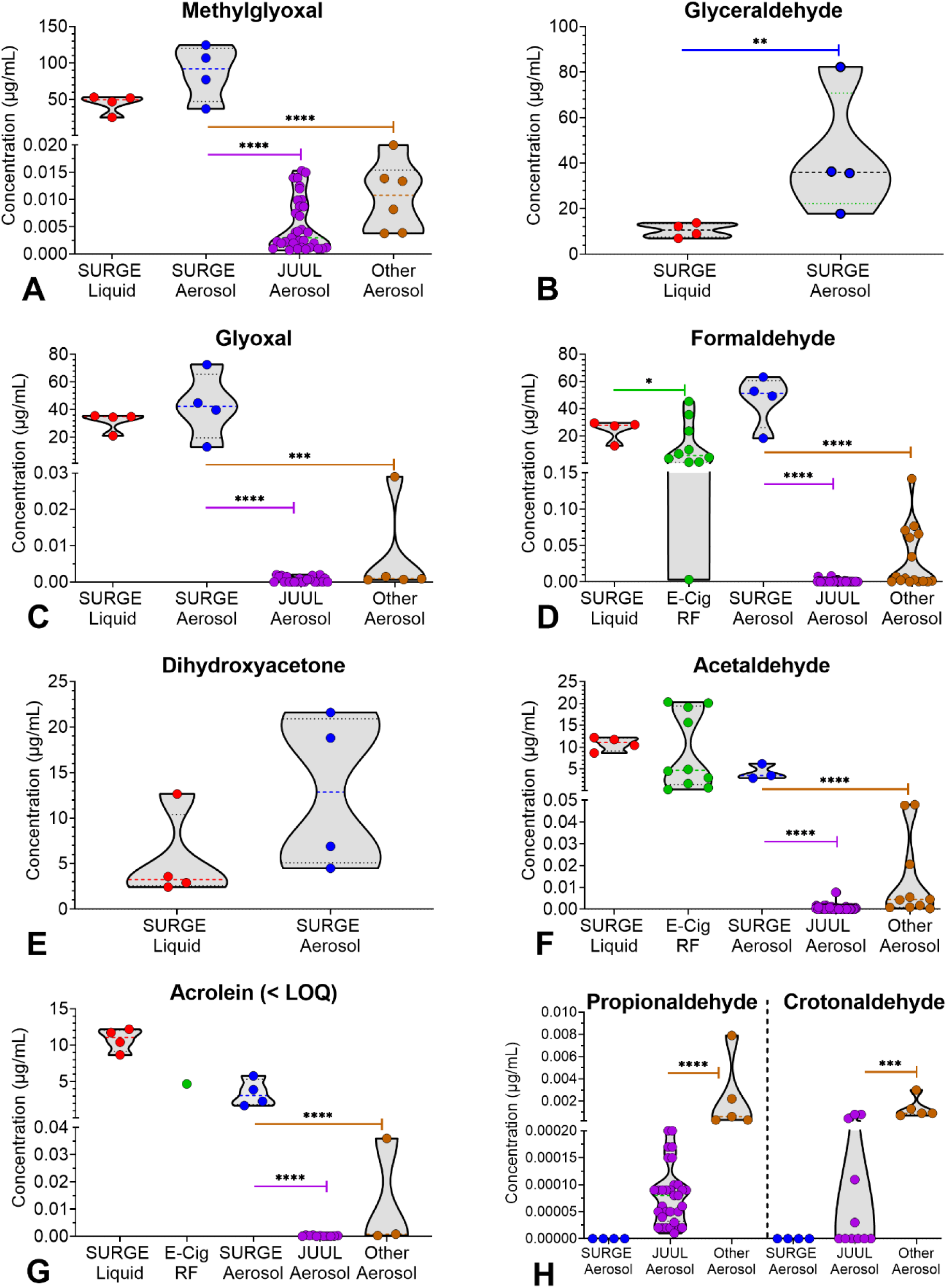
Concentrations of aldehydes in SURGE and e-cigarette products. (A) Methylglyoxal, (B) Glyoxal, (C) Glyceraldehyde, (D) Formaldehyde, (E) Dihydroxyacetone, (F) Acetaldehyde, and (G) Acrolein, (H) Propionaldehyde and Crotonaldehyde. E-Cig RF = e-cigarettes refill fluids. E-liquid data are taken from Gillman et al. 2020, Khlystov & Samburova 2016, LeBouf et al. 2018, Sleiman et al. 2016, Lim & Shin, 2013. E-cigarette aerosol data are taken from Azimi et al. 2021; Chen et al. 2021; Karam et al. 2022; Lalonde et al. 2022; Lestari et 2018; Mallock et al. 2020; Son et al. 2020; Talih et al. 2020; Talih et al. 2022; Uchiyama et al. 2013; Xu et al. 2023.

### Cytotoxicity of SURGE Fluids and Aerosols in the MTT Assay

The cytotoxicity of fluids and aerosols was evaluated with BEAS-2B cells using the MTT assay (Figures 5A-C). Products that reached an IC_70_ (30% lower value than the untreated control) were considered cytotoxic.^59^ All fluids were cytotoxic, with IC_70_s between 0.51– 0.59% and IC_50_s between 0.96 – 1.19% (Figure 5A and Table 1). “Watermelon Ice” and “Green Mint” aerosols captured in the culture medium were more cytotoxic than unvaped fluids (Figure 5B, Table 1). Total aldehyde concentrations in these two flavors were higher than in the other two products (Figure 3H).

**Figure 5.**
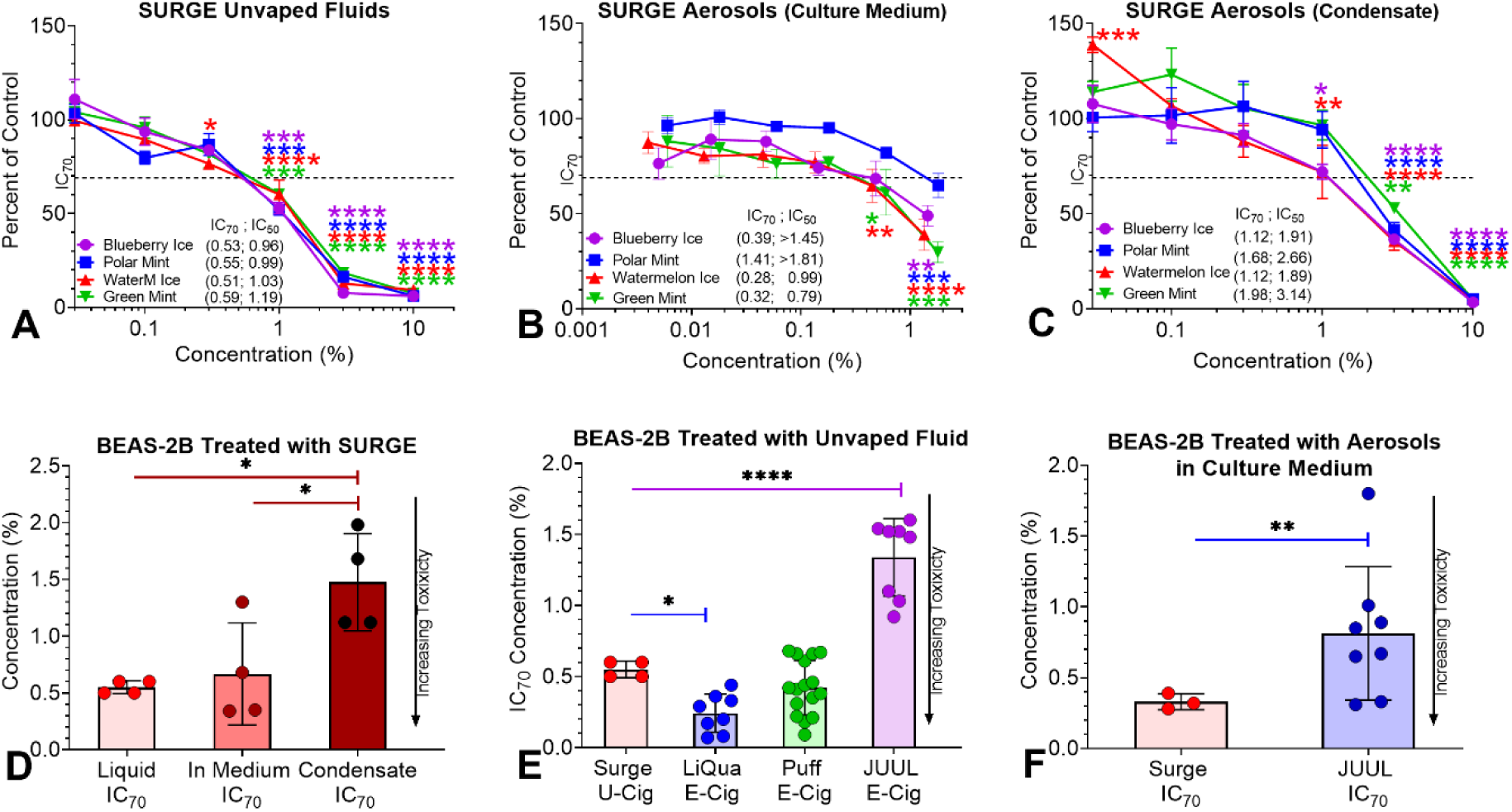
Concentration-response curves for BEAS-2B cells treated with SURGE products and Summary of MTT assay data for SURGE and e-cigarette products. (A) Unvaped SURGE fluids, (B) Surge aerosols captured in culture medium, (C) SURGE aerosols collected as condensates, (D) Comparison of SURGE fluids and aerosols. (E - F) Comparison of unvaped fluids and aerosol MTT IC_70_s for SURGE and e-cigarette products. Each point is the mean ± standard error of the mean of at least three independent experiments. U-Cig = u-cigarette, E-Cig = e-cigarettes. For statistical significance, * = p < 0.05, ** = p < 0.01, *** = p < 0.001, **** = p < 0.0001.

**Table 1.** IC_70_ and IC_50_ (%) of Surge U-Cigarette Pod Fluids and Aerosols in the MTT Assay.

| Surge Pod Flavors<sup>a</sup> | Unvaped Fluids |  | Aerosols in Medium |  |  | Aerosols Condensates |  |  |
| --- | --- | --- | --- | --- | --- | --- | --- | --- |
|  | IC<sub>70</sub> | IC<sub>50</sub> | IC<sub>70</sub> | IC<sub>50</sub> | Highest Conc. (%) | IC<sub>70</sub> | IC<sub>50</sub> | Highest Conc. (%) |
| Green Mint | 0.59 | 1.19 | 0.32 | 0.79 | 1.79 | 1.98 | 3.14 | 10 |
| Watermelon Ice | 0.51 | 1.03 | 0.28 | 0.99 | 1.35 | 1.12 | 1.89 | 10 |
| Blueberry Ice | 0.53 | 0.96 | 0.39 | > 1.45 | 1.45 | 1.12 | 1.91 | 10 |
| Polar Mint | 0.55 | 0.99 | 1.41 | > 1.81 | 1.81 | 1.68 | 2.66 | 10 |

a = order of pod flavors ranked according to IC<sub>70</sub> of aerosols made in medium from top to bottom.

For cells treated with aerosol condensates, both “Ice” flavors were slightly more toxic than the “Mint” flavors (Figure 5C). The SURGE aerosols collected in the culture medium were significantly more toxic than the aerosol condensates (Figure 5D).

The MTT cytotoxicity data (IC_70_s) of SURGE were compared to other pod-based e-cigarettes and LiQua refill fluids (Figures 5 E, F). For unvaped fluids, SURGE was significantly less cytotoxic than LiQua refill fluids,^39^ similar to PUFF fluids,^14^ and significantly more cytotoxic than JUUL fluids^13^ (Figure5E). For aerosols, SURGE was significantly more cytotoxic than JUUL (Figure 5F).

### Relationship between Cytotoxicity and Chemical Concentration

The relationship between the cytotoxicity observed with the fluids/aerosols in the MTT assay and chemical concentrations was determined using linear regression analysis (Figure 6A-I). Chemical concentrations in the ranges of 0.1 - 3% (unvaped fluids) and 0.013 – 1.8% (aerosol) were used to identify linear relationships. High (R^2^ ≥ 0.6) or moderate (R^2^ = 0.3 − 0.5) correlations were observed between cytotoxicity and the concentrations of propylene glycol, glycerol, nicotine, and WS-23 (Figure 6A-D). Correlation coefficients were slightly higher in the unvaped fluids than in the aerosols, with significant p-values (p <0.0001) for all fluid and aerosol treatments.

**Figure 6.**
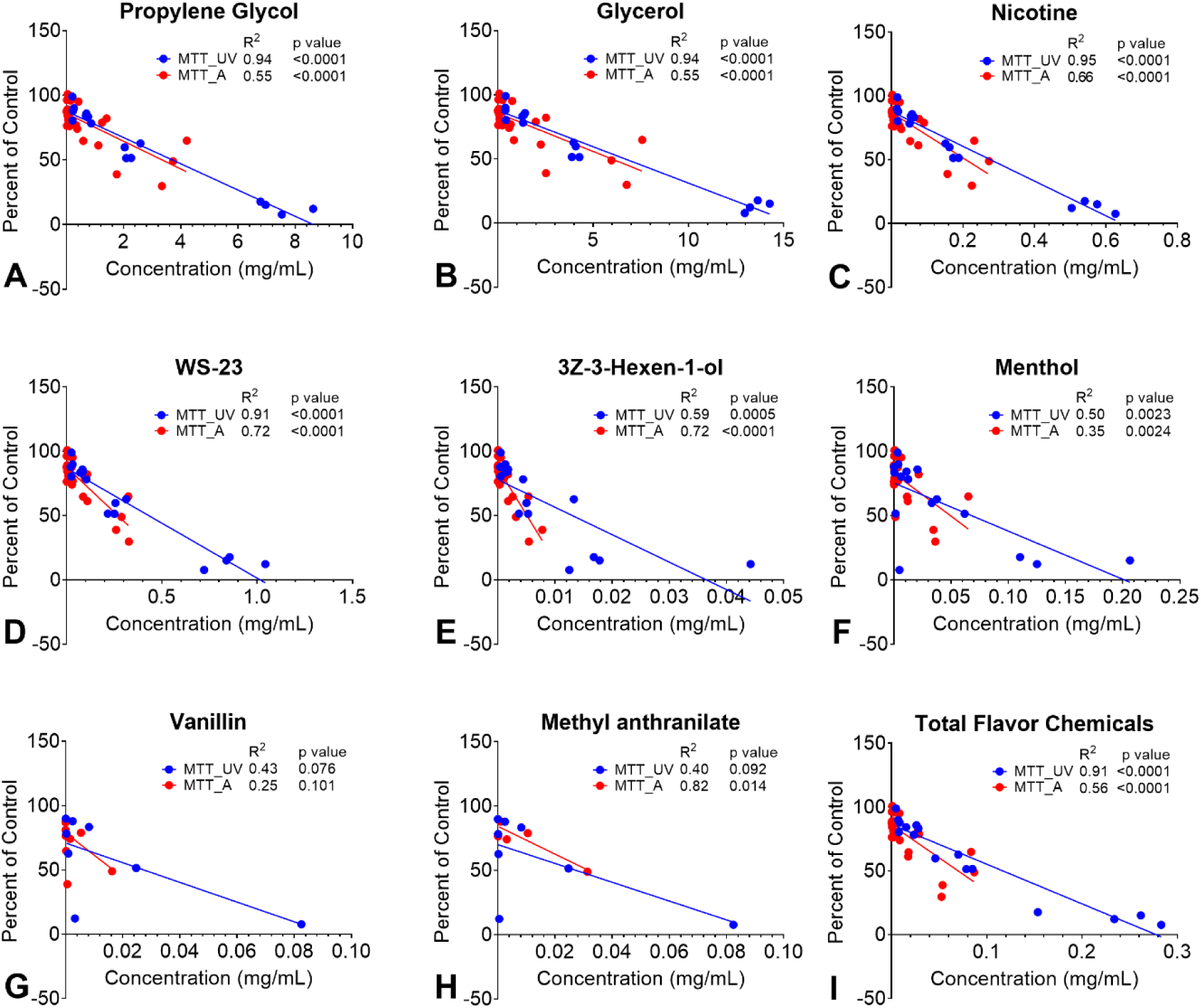
Relationship between cytotoxicity and dominant chemical concentrations. Regression analyses showing cytotoxicity on the *y*-axis, expressed as a percentage of the untreated control versus the concentrations of (A) Propylene Glycol, (B) Glycerol, (C) Nicotine, (D) WS-23, (E) 3Z-3-Hexen-1-ol, (F) Menthol, (G) Vanillin, (H) Methyl Anthranilate, and (I) Total Flavor Chemicals. Blue lines and dots are unvaped fluids (UV), and red lines and dots are aerosol (A).

Concentrations of dominant flavor chemicals were moderate to weakly correlated with toxicity except for (3Z)-3-hexen-1-ol and methyl anthranilate with aerosol R^2^s of 0.72 and 0.82, respectively. (Figure 6E - H). For total flavor chemicals, cytotoxicity was highly correlated (R^2^ = 0.91) for unvaped fluids and moderately correlated (R^2^ = 0.56) for aerosols. (Figure 6I).

### Relationship Between Cytotoxicity and Aldehyde Concentrations

**The** regression analyses for aldehydes and total aldehyde content in the fluids and aerosols showed strong correlations with significant p-values (Figure 7A-H). R^2^ was generally slightly higher for fluids than aerosols. Correlation coefficients of acetaldehyde, glyoxal, methylglyoxal, and formaldehyde were 0.79 - 0.93 with p < 0.0001 for fluid samples (Figure 7A-D). For acrolein, glyceraldehyde, and dihydroxyacetone with levels below the LOQ, correlation coefficients were moderate to high (0.4 – 0.93) with significant p values (Figure 7E-G). Cytotoxicity vs total aldehyde concentration had high R^2^s and significant p values (Figure 7H).

**Figure 7.**
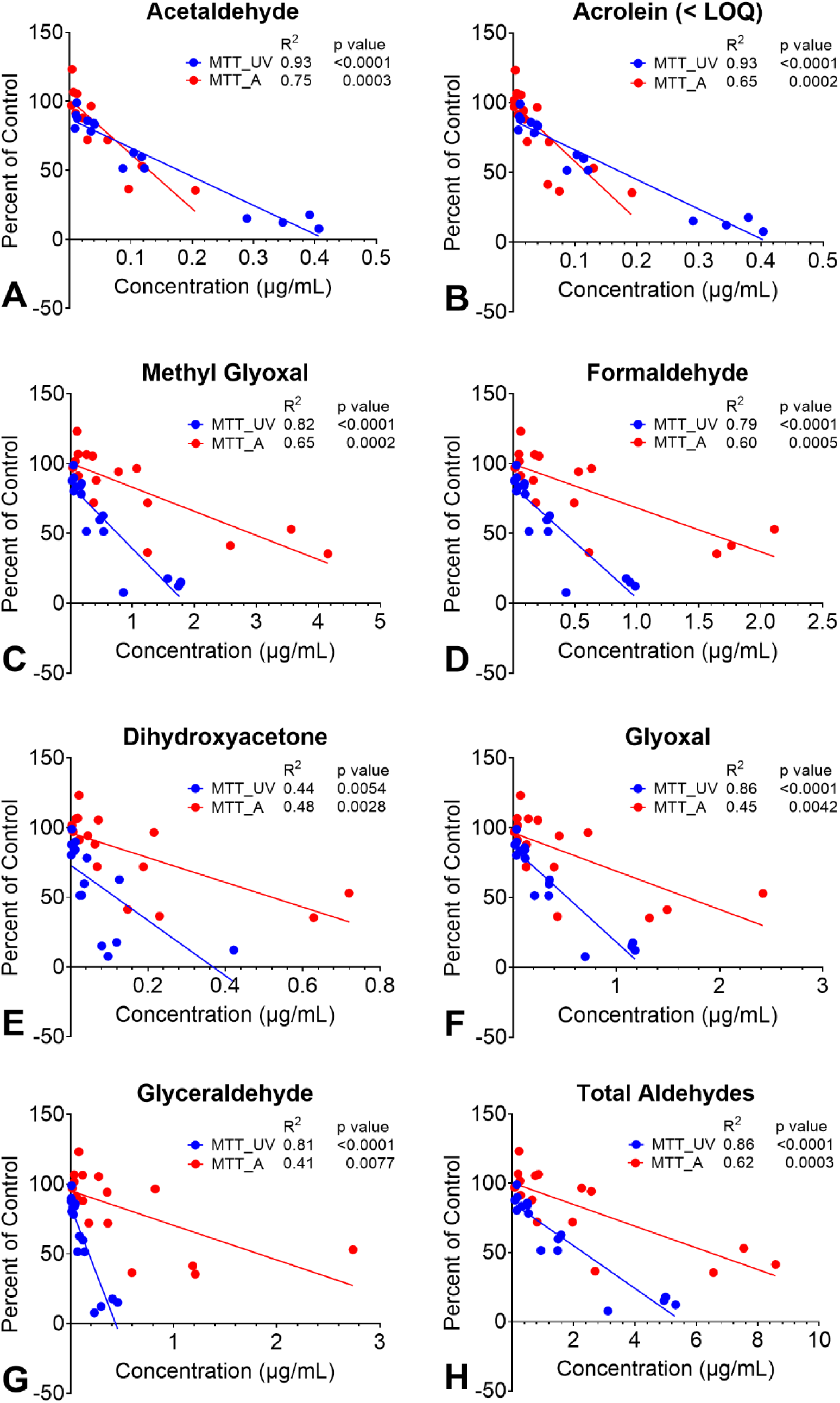
Relationship between cytotoxicity and aldehyde concentrations. Regression analyses showing cytotoxicity expressed as a percentage of the untreated control (y-axis) versus the concentrations of (A) Acetaldehyde, (B) Acrolein, (C) Methylglyoxal, (D) Formaldehyde, (E) Dihydroxyacetone, (F) Glyoxal, (G) Glyceraldehyde, and (H) Total Aldehydes. Blue lines and dots are unvaped fluids (UV), and red lines and dots are aerosols (A).

### SURGE Aerosols Inhibited Cell Growth and Altered Cell Morphology

In time-lapse videos, the area occupied by untreated control cells (0 TPE) increased throughout exposure (Figures 8 A-D), and cell morphology appeared normal, reaching confluency by 48 hours of incubation (Figures 8 E-H). However, cell area was significantly lower for BEAS-2B cells exposed to media containing 0.6 or 6.0 TPE of SURGE aerosols (Figures 8 A-D). Untreated control cells grew in a typical epithelial monolayer, while SURGE aerosol-treated cells were fewer and often attenuated or rounded, especially in the 6 TPE groups (Figure 6 E - H). At 6 TPE, “Polar Mint” and “Watermelon Ice” appeared to kill cells by 4 hours (Figures 8 F, H), while cells treated with “Blueberry Ice” and “Green Mint” were attenuated (Figures 8 E, G).

**Figure 8.**
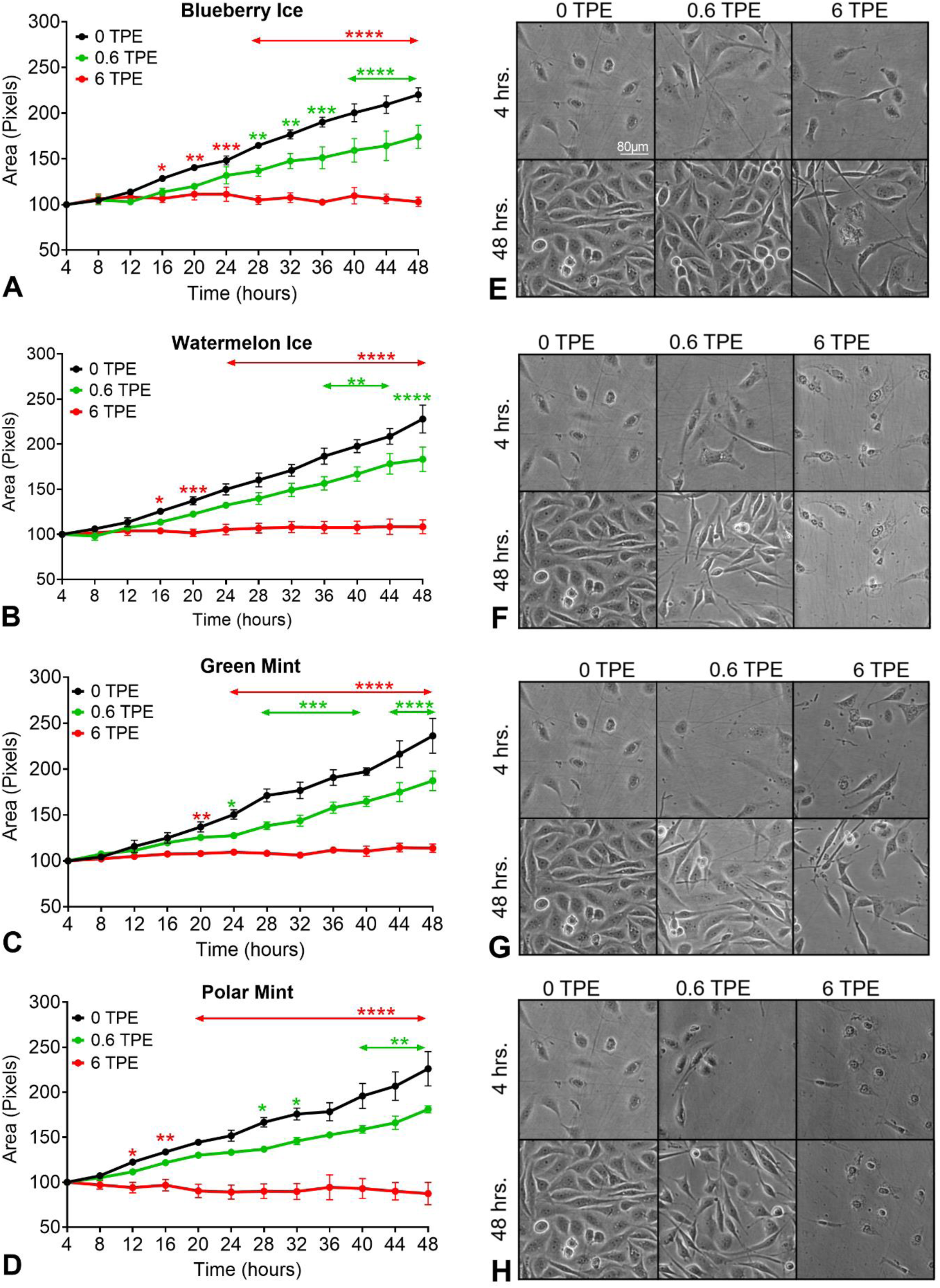
Effects of SURGE aerosols on cell growth (area) and morphology in the live-cell imaging assay. Time-lapse imaging was performed with (A) Blueberry Ice, (B) Watermelon Ice, (C) Green Mint, and (D) Polar Mint. The area covered by cells in the monolayer is shown over time. (E - H) Micrographs of BEAS-2B cells after 4 and 24 hours of treatment with 0, 0.6 TPE, or 6 TPE of SURGE aerosols. For A - D, each point is the mean of at least three experiments ± the SEM. * = p < 0.05; ** = p<0.01; *** = p < 0.001; **** = p<0.0001

### SURGE Aerosols Depolymerized Actin Filaments in BEAS-2B Cells

Based on the attenuated and rounded cell morphologies observed in the live cell imaging assay, the effect of aerosol treatment on actin filaments was examined at 0.6 and 6 TPE using phalloidin-TRITC, which labels f-actin (Figure 9). Untreated control cells were spread and had polymerized actin filaments in their cytosol and subjacent to their plasma membranes (Figure 9A white arrowheads). At 0.6 TPE, there was some evidence of actin filament depolymerization, but intact f-actin filaments were still present (Figure 9B white arrowheads). At 6 TPE, F-actin filaments were rarely observed underlying the plasma membrane and in the cytoplasm. Rather, small puncta of f-actin were present in the cytoplasm (yellow arrowheads), and f-actin extended into thin projections on the cell surface (blue arrowheads) (Figures 9 C-F). These projections were not present in the controls (Figure 9A).

**Figure 9.**
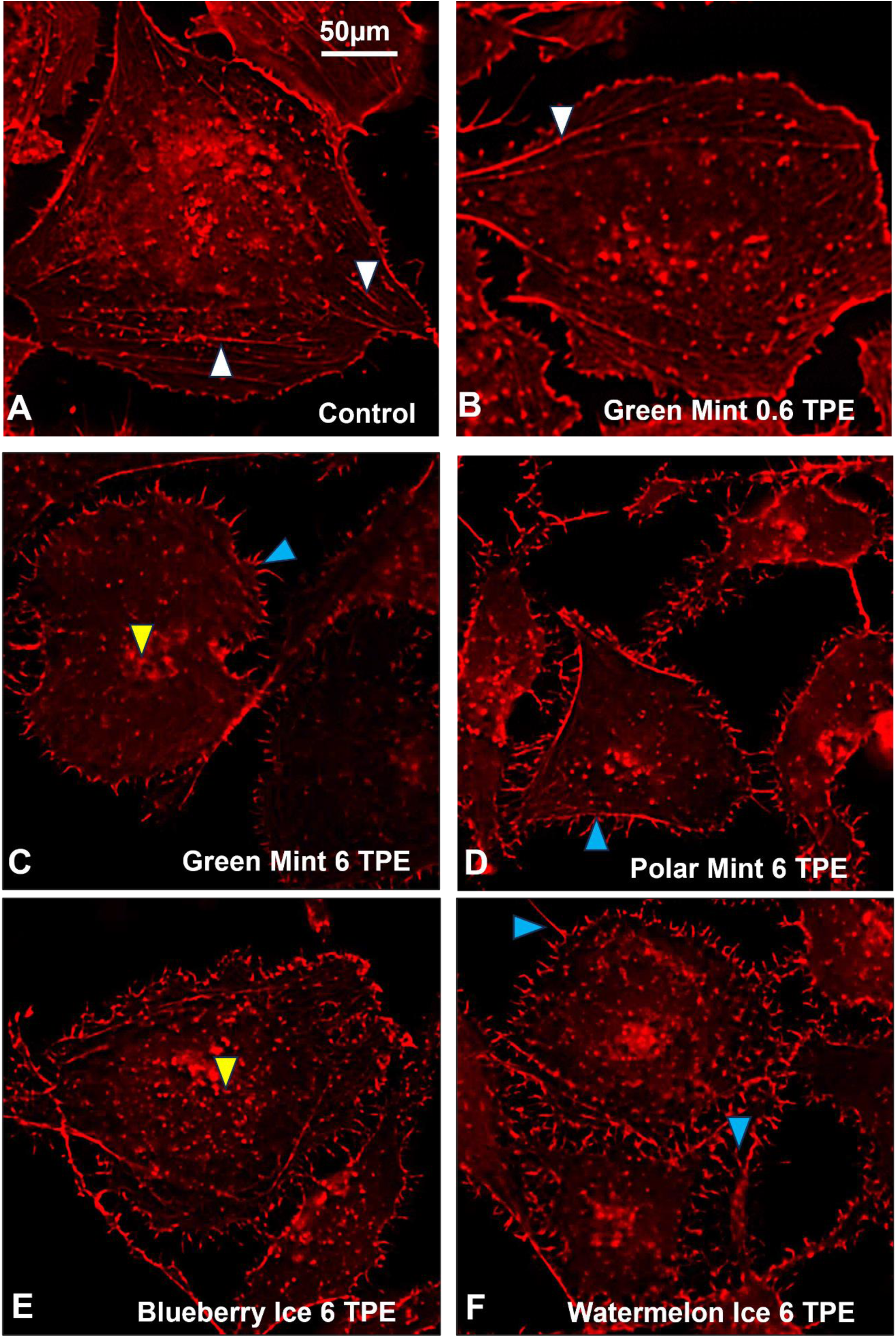
BEAS-2B cells treated with 0.6 or 6 TPE of SURGE aerosols. Treatment was for 24 hours, and then cells were labeled with phalloidin-TRITC. (A) Untreated control, (B - C) Green Mint, (D) Polar Mint, (E) Blueberry Ice, and (F) Watermelon Ice. White arrowheads in the control condition show f-actin in the cytoplasm and beneath the plasma membrane. In the SURGE treatment groups, blue arrowheads show f-actin in spikes on the cell surface, and yellow arrowheads show f-actin puncta in the cytoplasm.

### Margin of Exposure (MOE) Analysis for Chemicals in Unvaped SURGE Fluids

The MOE prioritizes the potential risk of exposure to chemicals and food additives.^60^ The potential risk of nicotine, WS-23, and propylene glycol from daily exposures was calculated using “no observed adverse effect levels” (NOAELs) from experimental data as a reference point,^61–63^ an estimated daily exposure to the chemical, and an average adult body weight of 60 kg. The MOE calculation was based on daily consumption of 1 or 2 SURGE /day (fluid volumes of 1.2 or 2.4 mL) and a 100% transfer from the fluid mixture into the aerosol. MOE values below the 100 threshold for a food additive and 1000 threshold for nicotine^62^ are considered high risk and require risk prioritization and mitigation by regulatory agencies. For nicotine, propylene glycol, and WS-23, all MOEs were <100 for all SURGE flavors at 1 or 2 pods consumption/day (Figure 10).

**Figure 10.**
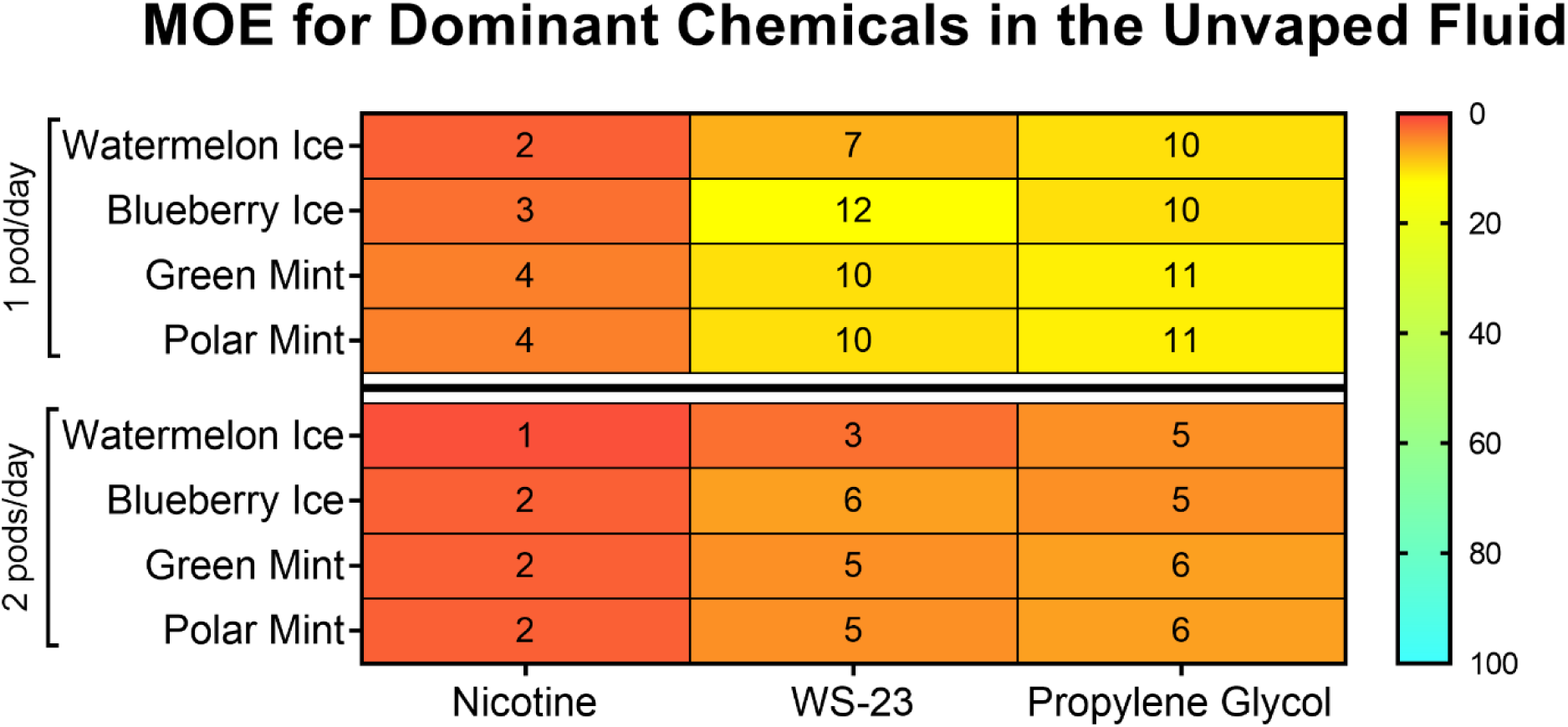
The margin of exposure (MOE) for nicotine, WS-23, and propylene glycol in SURGE u-cigarette fluids. MOEs are below the threshold of 1000 for nicotine and 100 for food additives, indicating a human health risk.

## DISCUSSION

The chemicals in SURGE fluids and aerosols were similar to those in multiple generations of e-cigarettes.^13,14,64–66^ The four flavored SURGE products had fluids and aerosols that contained high concentrations of propylene glycol, glycerol, nicotine, and flavor chemicals, in agreement with the SURGE website.^67^ However, the SURGE website does not mention the synthetic coolant WS-23, which is also used in high concentrations in Puff products.^14,16,17,68^ The concentrations of aldehydes in both SURGE fluids and aerosols were higher than in other pod-based products, and these were not mentioned on the SURGE website. Each SURGE flavor inhibited mitochondrial reductases in the MTT assay and cell growth in the live cell imaging assay. Toxicity in the MTT assay was higher than we have previously observed for JUUL and was highly correlated with the concentrations of propylene glycol, glycerol, nicotine, WS-23, and aldehydes. In treated groups, cell growth was inhibited dose-dependently, cell morphology was adversely affected in time-lapse images, and actin filaments were depolymerized in human bronchial epithelial cells. In MOE assessments, nicotine, WS-23, and propylene glycol concentrations in SURGE fluids were high enough to be a health concern based on consumption of 1 or 2 SURGE u-cigarettes/day. Collectively, these data question the safety of SURGE products.

While SURGE pods are labeled 18 mg/mL, the measured concentrations of nicotine ranged from 15 to 19 mg/L, with “Blueberry Ice” being slightly higher than the labeled value. Discrepancies between labeled and measured nicotine concentrations have also been reported for e-cigarettes and refill fluids.^69–71^ Nicotine concentrations in SURGE products were lower than in JUUL, Puff or other 4^th^ generation pod-based products but within the range of freebase nicotine previously reported in refill fluids^13,17,69,71–77^ suggesting SURGE does not use benzoic or other acids to create nicotine salts in their products. However, a negligible level of acetic acid, estimated at 60 µg/mL, was found in the “Blueberry Ice” flavor. Transfer of nicotine into the aerosol was between 74 and 100% efficient in SURGE and generally higher or similar to what has been reported in e-cigarettes.^13,74,78,79^ Nicotine concentrations in both fluids and aerosols were highly correlated with cytotoxicity in the MTT assay, as has also been reported for e-cigarettes.^13^

While WS-23 was present in SURGE fluids at concentrations within the range found in Puff fluids (range = 0.8 – 45 mg/mL),^14^ “Watermelon Ice” had one of the highest concentrations of WS-23 (31 mg/mL) we have seen in tobacco products. All WS-23 concentrations in SURGE were well above those recommended for use in cosmetics, hygiene, edibles, and household products (recommended range = 0.0008−0.3%).^80^ WS-23 transferred to the aerosol with good efficiency (62 to 92%), and like nicotine, its concentrations in both fluids and aerosols were highly correlated with toxicity in the MTT assay. WS-23 was originally developed by Wilkenson Sword for use in shaving cream^81,82^ and is a relatively new additive in e-cigarette products.^14,16,17^ It caused depolymerization of actin filaments in BEAS-2 B cells exposed at the air-liquid interface in a cloud chamber, leading to impaired motility and a collapse of cell architecture^83^ and may have been a factor in depolymerizing actin in cells treated with the SURGE aerosols.

Individual flavor chemicals were generally present in fluids at < 6 mg/mL, which is lower than most dominant flavor chemicals in JUUL and Puff products.^13,14^ Generally, the transfer of dominant flavor chemicals to SURGE aerosols ranged from 44% to 87%. The total flavor chemical concentrations in SURGE were within the range found in other pod e-cigarette brands (JUUL and Puff) and refill fluids (LiQua). ^13,14,39^ Individual and total flavor chemical concentrations generally correlated (R^2^ ≥ 0.5) with toxicity in the MTT assay.

An unexpected finding was the relatively high concentration of aldehydes in SURGE products. While fluid concentrations of aldehydes would be similar for all users and were high in SURGE compared to other 4th generation products, ^48–51,53–56,84^ numerous factors such as e-cigarette brand, solvent, heat, power, ingredients, user topography, and metals in atomizers can influence aldehyde concentrations in aerosols.^1,37,43,46,48,54,57,72,85,86^ For example, the average concentrations of methylglyoxal (45 µg/mL) and glyoxal (31 µg/mL) in the unvaped SURGE fluids were much higher than concentrations reported in e-cigarette aerosols (methylglyoxal = 0.006 µg/mL and glyoxal = 0.001 µg/mL).^48,50,54–56^ Even if no additional aldehydes formed during vaping, the baseline concentrations in unvaped fluids and their transfer to the aerosol during vaping would be a concern. The aldehydes may be introduced by individual ingredients when compounding the SURGE fluids, or they may form from interactions with other constituents in fluid mixtures.

Except for acetaldehyde in all products and glyoxal in “Blueberry Ice,” the concentrations of the aldehydes were higher in SURGE aerosols than in unvaped fluids. This increased concentration in aerosols is likely due to the formation of reaction products from propylene glycol and glycerol, as reported in coil-style e-cigarettes upon heating.^85,87^ The temperature range of SURGE products during ultrasonication is lower (120 – 150 °C) than in coil-style e-cigarettes (200 – 250 °C)^34^ leading SURGE to claim “Vapour is created at a lower working temperature, producing fewer potential “toxins” in SURGE aerosol than traditional coil-based pod-style e-cigarettes.”^34^ The logic of the statement on the SURGE website notwithstanding, we found higher concentrations of potential toxins in SURGE aerosol than in other pod-style e-cigarettes (Figure 4).

Aerosols collected as condensates were significantly less toxic than those collected in culture media, showing that the collection method affects endpoint data and that the condensation method was less efficient in preserving toxicity than the culture medium protocol. The SURGE website claims their products are less harmful than traditional coil-heated e-cigarettes.^34^ Nevertheless, unvaped fluids and aerosols from SURGE were significantly more toxic in the MTT assay than JUUL products^13^ (Figure 5). Toxicity in the MTT assay correlated most strongly with propylene glycol, glycerol, nicotine, WS-23, and total flavor chemicals, similar to correlations found for Puff fluids^14^ and JUUL aerosols and fluids.^13^ These data support the conclusion that SURGE fluids/aerosols are not less cytotoxic than fluids/aerosols produced by traditional e-cigarettes and that the chemicals in the highest concentrations contribute most to cytotoxicity. Aldehydes, which are well-established toxicants,^88^ also correlated well with cytotoxicity (R^2^ ≥ 0.6 for total aldehydes), even though their tested concentrations were lower than the dominant chemicals. Their presence in SURGE aerosols is also of concern since some, such as formaldehyde, are considered carcinogens.^89^

Aerosols from the four SURGE products inhibited BEAS-2B cell growth in a concentration-dependent manner, similar to Puff fluids.^14^ 6 TPE of “Polar Mint” and “Watermelon Ice” appeared to also cause cell death, which may have been due to the higher concentrations of dominant flavor chemicals and/or aldehydes in these flavors. At 6 TPE, all SURGE fluids caused depolymerization of actin filaments in BEAS-2B cells and the formation of actin-containing spikes at the cell surface. Cells with depolymerized actin were often flat and spread, suggesting that their microtubules and/or intermediate filaments were still intact. Pure WS-23 caused depolymerization of actin filaments in an air-liquid interface exposed 3D bronchial epithelium and monolayers of BEAS-2B cells.^83^ and may have been responsible for the depolymerization of actin caused by SURGE. Similar to prior data with 16 Puff products,^14^ the concentrations of WS-23 in SURGE unvaped fluid were high enough to produce MOEs below the threshold of 100, indicating a potential health risk. In SURGE, the other dominant chemicals (nicotine and propylene glycol) also had MOEs below 100. Concerns about the high concentrations of nicotine, solvents, and flavor chemicals have been raised previously with e-cigarettes^13,90^ and remain a concern with u-cigarettes.

Although the idea that u-cigarettes are safer than those with heated coils appears logical, it is not supported by SURGE cytotoxicity data, which show that SURGE aerosols are as toxic or more toxic than the JUUL and Puff comparators and support the conclusion that high concentrations of the dominant chemicals in SURGE and the relatively high concentrations of aldehydes are a source of toxicity. The SURGE website states that the particle size in their aerosols is smaller than other brands, improving the taste of their product.^36^ Small particles would also penetrate deeper into the lungs,^91^ which may not be desirable.^92^

### Limitations

The concentration of aldehydes in SURGE aerosols could vary depending on user topography, which is highly variable^37^ and could be higher for some users than those reported here. Moreover, our aldehyde concentrations in aerosols are based on condensates which were less toxic than aerosols collected in culture media and may, therefore, underestimate the actual concentrations users receive. We examined only one brand of u-cigarette, and others should be evaluated in the future.

In conclusion, unvaped SURGE liquids are similar in basic composition to other 4th generation pod-style e-cigarette fluids but appear to use freebase nicotine rather than nicotine salt, found in JUUL and Puff products.^13,14,16,17,53,74^ WS-23 was in SURGE products at concentrations well above those recommended for consumer products. Aldehyde concentrations were unusually high in unvaped fluids and generally higher in SURGE aerosols. Transfer of chemicals, including aldehydes, to aerosols generally occurred with good to excellent efficiency. SURGE cytotoxicity was highly correlated with concentrations of nicotine, WS-23, total flavor chemicals, and total aldehydes. SURGE aerosols also inhibited cell growth, similar to other 4th generation pod products. SURGE fluids depolymerized actin filaments, perhaps due to the high concentration of WS-23 in SURGE products. While the manufacturer claims that “SURGE produces the purest vapor of any device on the market,” data suggest that SURGE aerosols are very similar to other 4th generation aerosols and are higher in aldehydes known to be toxicants than other 4th generation pod products. SURGE is a good example of why new products must be tested before marketing since logical ideas (“lower heat produces fewer potential toxins”)^34^ do not necessarily hold up when tested. The elevated levels of aldehydes in SURGE products are a health concern, and users should be cautious and not assume they are safer than other pod-style e-cigarettes.

## Author Contributions

PT, JFP, and EO formed the concept and design of this study. EO, WL, and KJM carried out sample preparation, data collection, processing, and analysis. EO and PT wrote the first draft of the manuscript. EO, WL, KJM, JFP, and PT edited the manuscript.

## ACKNOWLEDGMENT

We want to thank Erik Ramirez, Onur Gazioglu, and Samantha Vargas for their help in making the aerosols and the UCR Stem Cell Core, which provided access to some of the instrumentation used in this project.

