## Supplementary material for "Ultrasonic Cigarettes: Chemicals and Cytotoxicity are Similar to Heated-Coil Pod-Style Electronic Cigarettes": Figure S1

### Ultrasonic Cigarettes (U-Cigarettes): Design, Chemical Analysis, and Cellular Effects of Fluids and Aerosols

[†]Department of Molecular, Cell, and Systems Biology. University of California, Riverside, California 92521, USA

[‡]Department of Civil and Environmental Engineering, Portland State University, Portland, Oregon 97207, USA

[§]Department of Chemistry, Portland State University. Portland, Oregon 97207, USA

**Corresponding Author**
*

**TABLE OF CONTENTS**

The Supporting Information is available free of charge.

PAGE S3 Heat map of flavor chemicals above the LOQ with concentrations below 1 mg/mL

PAGE S4 Flavor Chemicals Detected Below the Limit of Quantification

PAGE S5 Non-target Chemicals in U-Cigarettes

PAGE S6 List of target aldehydes

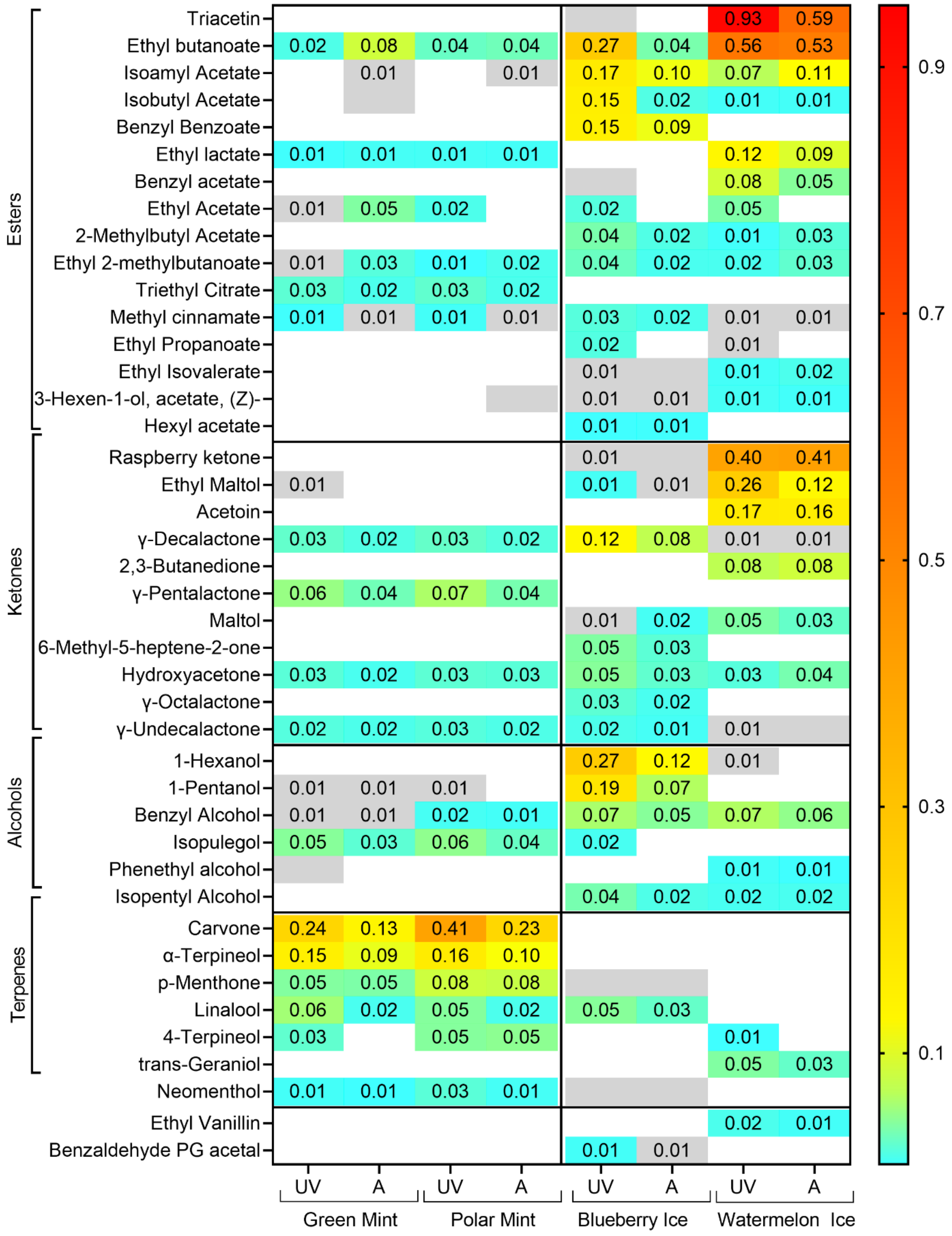


**Supplementary Figure 1.** Heat map of flavor chemicals above the LOQ (0.02 mg/mL) with concentrations below 1 mg/mL in u-cigarettes. Chemicals are ordered on the *y*-axis based on chemical class and frequency of occurrence of flavor chemicals from top to bottom within each class. Products are grouped on the *x*-axis according to flavor type (mint vs ice). The color gradient on the right shows the concentrations of the flavor chemicals in the heat map. Grey spaces indicate concentrations <LOQ.

**Supplemental Table 1. Flavor Chemicals Below the Limit of Quantification ((10 ug/mL for Unvaped Fluids and 20 µg/mL for Aerosols)**

| | Compound Name | CAS Number | Green Mint | | Polar Mint | | Blueberry Ice | | Watermelon Ice | | Chemical Class | Hazards[1] Class | Odor Type |
|---|---|---|---|---|---|---|---|---|---|---|---|---|---|
| | | | UV | A | UV | A | UV | A | UV | A | | | |
| 1 | Isoamyl Butyrate | 106-27-4 | 4.6 | 13.1 | 5.4 | 13.3 | ND | ND | 2.1 | 4.4 | Ester | Irritant | fruity |
| 2 | Menthyl Acetate | 16409-45-3 | 3.2 | 5.7 | 6.5 | 8.7 | 2.5 | 4.8 | 8.9 | 11.8 | Ester | ND | mentholic |
| 3 | Limonene | 138-86-3 | ND | 9.4 | ND | 10.7 | ND | ND | ND | ND | Terpene | Irritant | floral |
| 4 | Strawberry Glycidate_A | 77-83-8 | 4.5 | ND | ND | ND | 6.1 | 0 | 9.1 | 7.1 | Ester | ND | ND |
| 5 | Fenchol | 1632-73-1 | 5.8 | ND | 9 | ND | ND | ND | ND | ND | Terpene | Irritant | camphoreous |
| 6 | Amyl Acetate | 628-63-7 | 3.4 | 6 | 3.3 | ND | ND | ND | 8.8 | 5.1 | Ester | Irritant | fruity |
| 7 | β-Damascone | 85949-43-5 | 5.1 | 4.4 | 8.2 | 5.2 | 7.5 | 4.6 | 5.1 | 4 | Ketone | Irritant | fruity |
| 8 | Ethyl Cinnamate | 103-36-6 | ND | ND | ND | ND | ND | ND | 8.2 | 5.9 | Ester | Irritant | Balsamic |
| 9 | 2-Hexen-1-ol, (E)- | 928-95-0 | ND | ND | ND | ND | ND | ND | 8.2 | 3.5 | Alcohol | Irritant | fruity |
| 10 | α-Ionone | 127-41-3 | ND | ND | ND | ND | 8.2 | 6.7 | ND | ND | Ketone | Irritant | floral |
| 11 | Corylone | 765-70-8 | 6.1 | 2.4 | 7.9 | 3.1 | ND | ND | ND | ND | Ketone | Harmful | caramellic |
| 12 | Styralyl Acetate | 93-92-5 | 3.6 | ND | 4.6 | ND | 7.8 | 7.4 | ND | ND | Ester | ND | green |
| 13 | δ-Dodecalactone | 713-95-1 | ND | ND | ND | ND | ND | ND | 6.2 | 5.5 | Ketone | Irritant | Tropical |
| 14 | Amyl Isovalerate | 25415-62-7 | ND | ND | ND | ND | ND | ND | 7.5 | ND | Ester | ND | fruity |
| 15 | 2,3,5,6-Tetramethylpyrazine | 1124-11-4 | ND | ND | ND | ND | ND | ND | 7.4 | ND | Pyrazine | Harmful | nutty |
| 16 | Benzyl DMC butyrate | 10094-34-5 | ND | ND | ND | ND | 6.5 | 7 | ND | ND | Ester | Irritant | floral |
| 17 | Acetylpyrazine | 22047-25-2 | 4.6 | ND | 6.6 | 3.4 | ND | ND | ND | ND | Pyrazine | Irritant | popcorn |
| 18 | Pulegone | 89-82-7 | 4.5 | ND | 6.6 | 3.5 | ND | ND | ND | ND | Terpene | Harmful | minty |
| 19 | (E)-β-Ionone | 79-77-6 | ND | ND | ND | ND | 5.4 | 5 | 1.6 | ND | Ketone | Irritant, D | floral |
| 20 | Piperitone | 89-81-6 | 3.1 | 1.8 | 5.1 | ND | ND | ND | ND | ND | Ketone | Irritant | herbal |
| 21 | 2,3,5-Trimethylpyrazine | 14667-55-1 | 4.4 | ND | 4.5 | 3.1 | ND | ND | ND | ND | Pyrazine | Harmful | nutty |
| 22 | β-Pinene | 127-91-3 | ND | 3.5 | ND | 4.3 | ND | ND | ND | ND | Terpene | Harmful, D | herbal |
| 23 | Benzaldehyde | 100-52-7 | ND | ND | ND | ND | 0.9 | ND | 4.1 | 3.1 | Aldehyde | Harmful | fruity |
| 24 | Acetophenone | 98-86-2 | ND | ND | ND | ND | ND | ND | 3.8 | 3.5 | Ketone | Harmful | floral |
| 25 | Strawberry Glycidate_B | 77-83-8 | ND | ND | ND | ND | ND | ND | 2.4 | ND | Ester | ND | ND |
| 26 | α-Pinene | 80-56-8 | ND | 2.5 | ND | 2.6 | ND | ND | ND | ND | Terpene | Harmful, D | herbal |
| 27 | Guaiacol | 90-05-1 | ND | ND | ND | ND | ND | 2 | ND | ND | Phenol | Harmful | Phenolic |
| 28 | Thymol | 89-83-8 | 1 | ND | ND | ND | ND | ND | ND | ND | Phenol | Corrosive, D | herbal |

[1]D = Dangerous to the environment. ND = Not Determined. UV = unvaped. V = vaped

**Supplemental Table 2. Non-target Chemicals in U-Cigarettes**

| Sample | Non-targets | Minor Non-target |
|---|---|---|
| Blueberry Ice | 1-ACETOXY-2-PROPANOL | Likely PG acetate ester isomer |
| | 1,2-PROPANEDIOL, 2-ACETATE | |
| | Octamethyltetrasiloxane | Siloxane from silicone rubber etc. |
| | 1,2,3-Propanetriol, monoacetate | GL acetate ester (monoacetin) |
| | Diacetin | GL acetate ester |
| | Raspberry ketone PG KETAL | |
| | 2,2,4,4,6,6,8,8-Octamethyl-1,3,5,7,2,4,6,8-Tetraoxatetrasilocane | Siloxane from silicone rubber |
| Watermelon Ice | 1-Acetoxy-2-Propanol | Likely PG acetate ester isomer |
| | Octamethylcyclotetrasiloxane | Siloxane from silicone rubber etc. |
| | 2,6-Dimethylhept-5-Enal | |
| | Methyl Dihydrojasmonate | |
| | 4-Octylbutan-4-Olide | |
| Polar Mint | 1-Acetoxy-2-Propanol | Likely PG acetate ester isomer |
| | 2-Acetoxy-1-Propanol | Likely PG acetate ester isomer |
| | Octamethylcyclotetrasiloxane | Siloxane from silicone rubber |
| | 1,2,3-Propanetriol, monoacetate | Likely GL acetate ester isomer |
| | (-)-Beta-Fenchol | |
| Polar Mint | 1-Acetoxy-2-Propanol | Likely PG acetate ester isomer |
| | 2-Acetoxy-1-Propanol | Likely PG acetate ester isomer |
| | Octamethylcyclotetrasiloxane | Siloxane from silicone rubber |
| | 1,2,3-Propanetriol, monoacetate | GL acetate ester (monoacetin) |

Acetate esters of propylene glycol and glycerol and siloxanes from silicon rubber were non-target chemicals estimated in u-cigarette aerosols (Table S2).

**Table 2. List of target aldehydes**

| # | Compound Name |
|---|---|
| 1 | Formaldehyde |
| 2 | Acetaldehyde |
| 3 | Acrolein |
| 4 | Propanal |
| 5 | Butanal |
| 6 | Crotonaldehyde |
| 7 | Pentanal |
| 8 | Glyceraldehyde |
| 9 | Dihydroxyacetone |
| 10 | 5-Hydroxymethylfurfural |
| 11 | Glyoxal |
| 12 | Methyl glyoxal |
